# Biotransformation and biodefluorination of chlorinated polyfluorocarboxylic acids by *Acetobacterium* species

**DOI:** 10.64898/2026.01.23.701208

**Authors:** Shun Che, Yaochun Yu, Weiyang Zhao, Yujie Men

**Author notes:** **Corresponding Author** Shun Che, Yujie Men.

## Abstract

Chlorinated per- and polyfluoroalkyl substances (Cl-PFAS) represent an important subgroup in replacement chemicals for legacy PFAS, and they have been detected in various environments including water bodies, soil, as well as air particles. However, the biodegradability of these chemicals is largely unknown. A recent study reported dechlorination triggered anaerobic biodefluorination of Cl-PFCAs by activated sludge communities. In this study, we further investigated the biodefluorination of Cl-PFCAs by pure cultures. Six of ten selected Cl-PFCAs displayed significant defluorination by *Acetobacterium bakii*, and corresponding pathways were proposed according to transformation product analysis. Several additional *Acetobacterium* species were all capable of defluorinating 3,5,7,8-tetrachloro-2,2,3,4,4,5,6,6,7,8,8- undecafluorooctanoic acid (CTFE4) with >50% removal of organic fluorine. On the contrary, *Clostridium homopropionicum* cannot defluorinate CTFE4. Crude protein extraction experiment showed that the enzymes responsible for CTFE4 dechlorination and defluorination required anaerobic atmosphere to function. Differential gene expression was analyzed according to RNA sequencing analysis, and the results indicated vitamin B12 related enzymes may involve in CTFE4 dechlorination.

## 1. Introduction

Per- and polyfluoroalkyl substances (PFAS), a huge group of man-made chemicals containing over 8,000 individual structures, have raised great concern on public health and ecosystems due to their persistence to degradation, bioaccumulation potential, and toxicity to human beings and ecosystems.^1, 2^ PFAS are widely used in manufacturing as well as everyday consumer products including fast food packaging, water resistant cloth, surfactants, pesticide, and firefighting foam, etc.^3^ The strong carbon-fluorine bonds endow PFAS with great stability to environmental degradation, and thus can remain in environments and human bodies for years.^4^ The potential health risk of exposure to PFAS includes cancer, liver damage, decreased fertility, and increased risk of asthma and thyroid disease, etc.^5^ Some legacy PFAS are now under regulation due to their potential threats on public health and ecosystem. The two most heavily used legacy PFAS, PFOA and PFOS, were phased out in North America and Europe. Due to the regulatory pressure, manufacturers have been switching to replace legacy PFAS with alternative fluorinated compounds, which have different functional groups such as fluoroalkylether, carbon-carbon double bonds, and hydrogen- or chlorine-substitutions.^6, 7^ However, a few replacement chemicals such as 6:2 Cl-PFESA and HFPO-DA were reported to have similar or even higher toxicity than legacy PFAS.^8^ Previous research mostly focused on legacy PFAS, while limited information of fate, transport, and toxicity has been obtained for the alternative fluorochemicals with fast-growing number of structures.

Cl-PFAS is one important subgroup in alternative fluorinated compounds for legacy PFAS, and they could be generated from different sources.^9^ For example, Cl-PFAS are used as building blocks for polychlorotrifluoroethylene (PCTFE), which is a widely used fluoropolymer.^8^ Historical manufacturing in fluorine industry can also emit Cl-PFAS as by products.^1^ According to functional group, Cl-PFAS could be further classified into several categories including chlorinated per- and polyfluoroalkyl carboxylic (Cl-PFCA), sulfonic acids (Cl-PFSA), ether acids and alcohols, as well as chlorofluoropolymer oligomers, etc. Detection of these Cl-PFAS have been emerging in last decade in various environments including waterbodies, air particles, as well as soils.^10-12^ However, their fate, transport, and biodegradability are largely unknown.

Few studies reported biotransformation and biodefluorination for Cl-PFAS.^13-15^ Compared to the strong C–F bonds, C–Cl bonds are more readily to be microbially cleaved due to lower bond energy.^15^ Reductive dechlorination of 6:2 and 8:2 chlorinated polyfluoroalkyl ether sulfonate (Cl-6:2 PFESA and Cl-8:2 PFESA) were observed in sediment environment and animals, forming H-6:2 PFESA and H-8:2 PFESA, respectively.^16, 17^ Although no defluorination was reported in these studies, dechlorination of Cl-PFAS may introduce delicate position for initiating microbial defluorination. A recent study reported that Cl-PFCAs could be biodefluorinated by activated sludge communities under anaerobic condition, and fluorine removal was triggered by dechlorination, identifying corresponding transformation products with HR-LCMS.^18^ Inspired by this finding, we further investigate the biodefluorination of Cl-PFCAs with pure cultures.

In this study, we aimed to understand the defluorination of different Cl-PFCAs by pure strains instead of complex microbial communities and explore the defluorination mechanisms. Biodefluorination of Cl-PFCAs were tested with different *Acetobacterium* species, and defluorination pathways were proposed according to transformation products analysis and were compared with those by activated sludge community. Defluorination mechanisms were explored by crude protein extraction and RNA expression analysis.

## 2. Materials and Methods

### 2.1 Chemicals

The ten chlorinated polyfluorocarboxylic acids **(Figure 1A** and details in **Table 1 & Figure S1**) were purchased from SynQuest Labs (Alachua, FL).

**Table 1.**
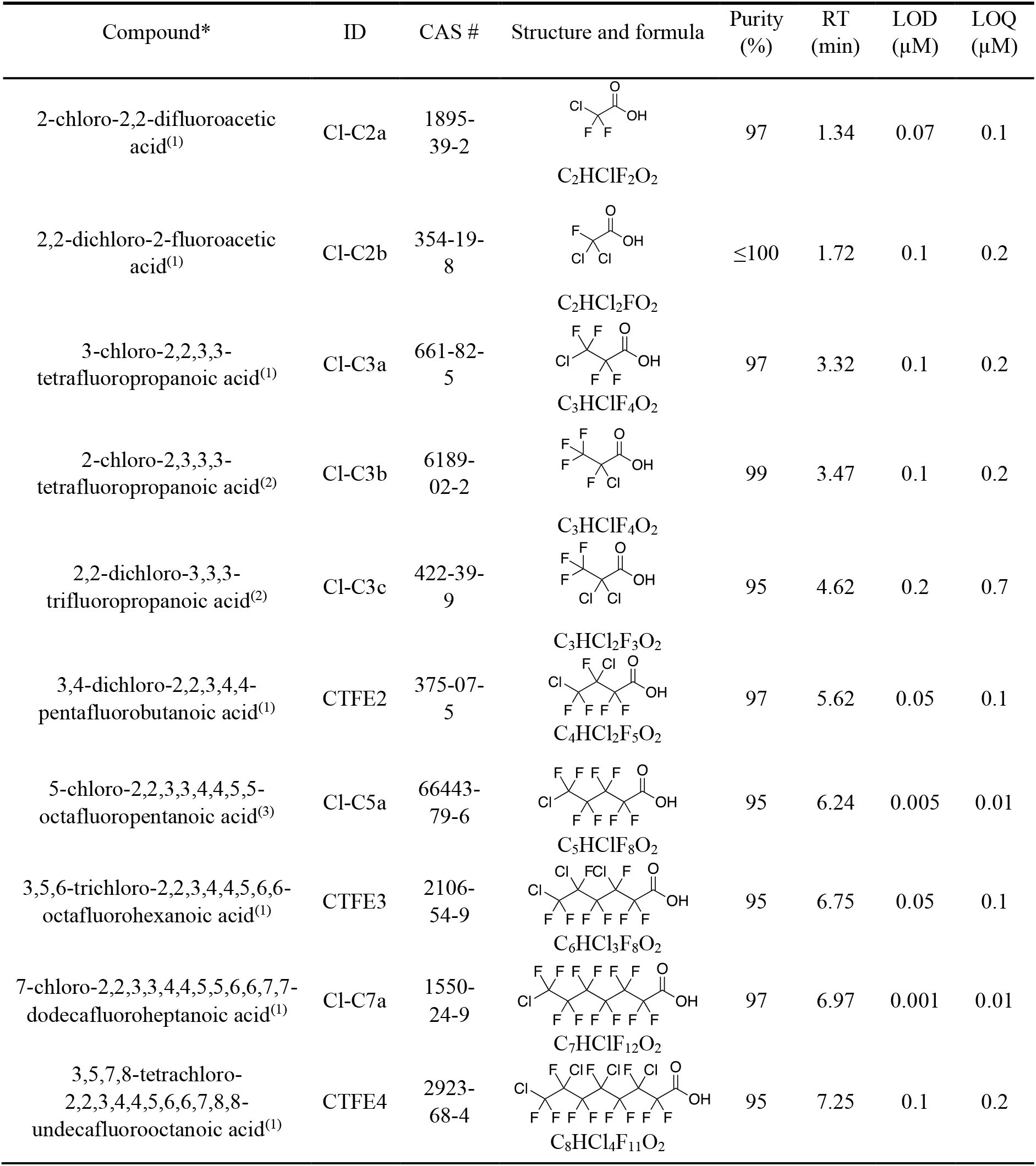
Cl-PFCAs screened for biodefluorination by *A. bakii*.

| Compound* | ID | CAS # | Structure and formula | Purity (%) | RT (min) | LOD (μM) | LOQ (μM) |
| --- | --- | --- | --- | --- | --- | --- | --- |
| 2-chloro-2,2-difluoroacetic acid<sup>(1)</sup> | Cl-C2a | 1895-39-2 | C<sub>2</sub>HClF<sub>2</sub>O<sub>2</sub> | 97 | 1.34 | 0.07 | 0.1 |
| 2,2-dichloro-2-fluoroacetic acid<sup>(1)</sup> | Cl-C2b | 354-19-8 | C<sub>2</sub>HCl<sub>2</sub>FO<sub>2</sub> | ≤100 | 1.72 | 0.1 | 0.2 |
| 3-chloro-2,2,3,3-tetrafluoropropanoic acid<sup>(1)</sup> | Cl-C3a | 661-82-5 | C<sub>3</sub>HClF<sub>4</sub>O<sub>2</sub> | 97 | 3.32 | 0.1 | 0.2 |
| 2-chloro-2,3,3,3-tetrafluoropropanoic acid<sup>(2)</sup> | Cl-C3b | 6189-02-2 | C<sub>3</sub>HClF<sub>4</sub>O<sub>2</sub> | 99 | 3.47 | 0.1 | 0.2 |
| 2,2-dichloro-3,3,3-trifluoropropanoic acid<sup>(2)</sup> | Cl-C3c | 422-39-9 | C<sub>3</sub>HCl<sub>2</sub>F<sub>3</sub>O<sub>2</sub> | 95 | 4.62 | 0.2 | 0.7 |
| 3,4-dichloro-2,2,3,4,4-pentafluorobutanoic acid<sup>(1)</sup> | CTFE2 | 375-07-5 | C<sub>4</sub>HCl<sub>2</sub>F<sub>5</sub>O<sub>2</sub> | 97 | 5.62 | 0.05 | 0.1 |
| 5-chloro-2,2,3,3,4,4,5,5-octafluoropentanoic acid<sup>(3)</sup> | Cl-C5a | 66443-79-6 | C<sub>5</sub>HClF<sub>8</sub>O<sub>2</sub> | 95 | 6.24 | 0.005 | 0.01 |
| 3,5,6-trichloro-2,2,3,4,4,5,6,6-octafluorohexanoic acid<sup>(1)</sup> | CTFE3 | 2106-54-9 | C<sub>6</sub>HCl<sub>3</sub>F<sub>8</sub>O<sub>2</sub> | 95 | 6.75 | 0.05 | 0.1 |
| 7-chloro-2,2,3,3,4,4,5,5,6,6,7,7-dodecafluoroheptanoic acid<sup>(1)</sup> | Cl-C7a | 1550-24-9 | C<sub>7</sub>HClF<sub>12</sub>O<sub>2</sub> | 97 | 6.97 | 0.001 | 0.01 |
| 3,5,7,8-tetrachloro-2,2,3,4,4,5,6,6,7,8,8-undecafluorooctanoic acid<sup>(1)</sup> | CTFE4 | 2923-68-4 | C<sub>8</sub>HCl<sub>4</sub>F<sub>11</sub>O<sub>2</sub> | 95 | 7.25 | 0.1 | 0.2 |

**Figure 1.**
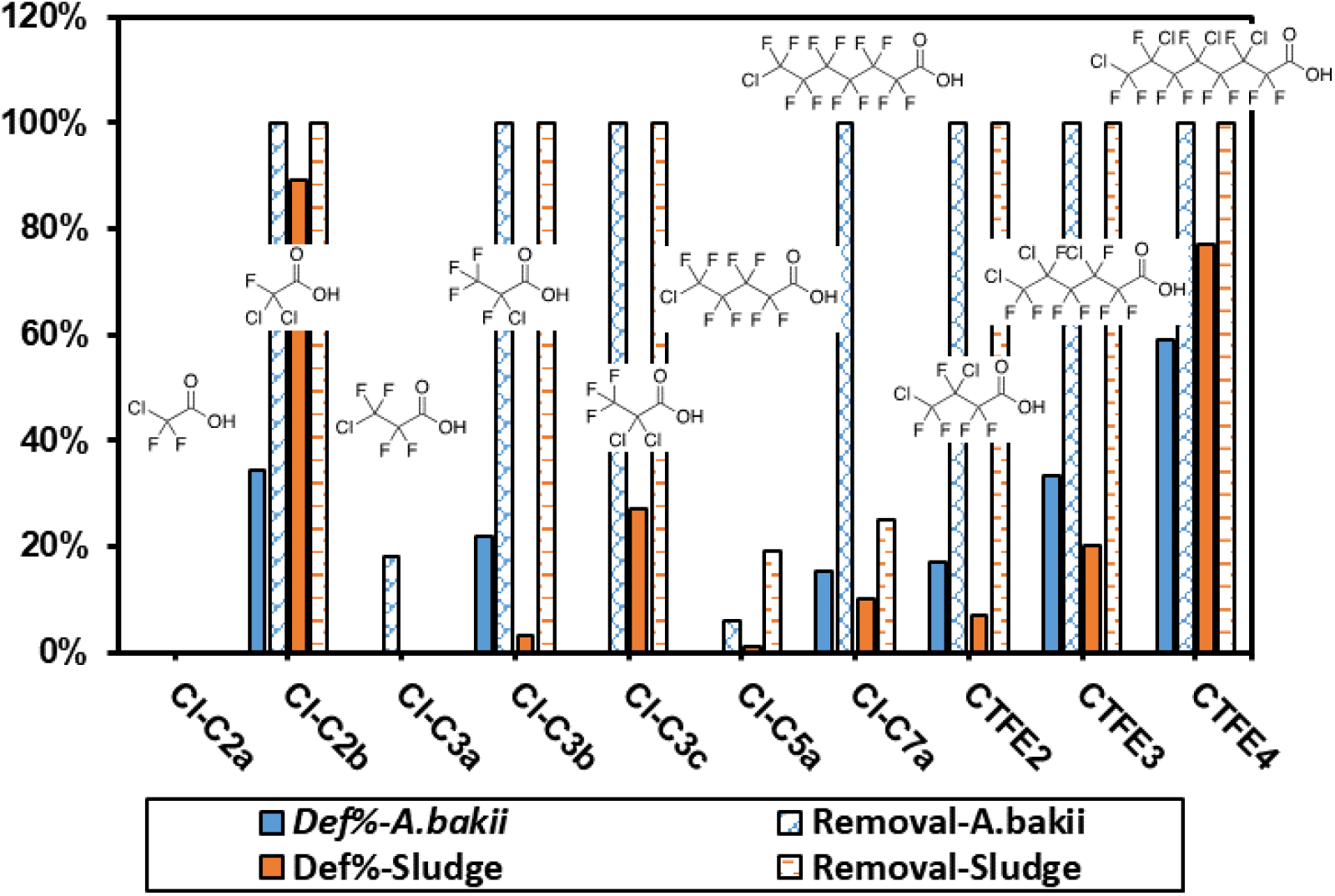
Defluorination degree of Cl-PFCAs in *A. bakii* and sludge communities, data acquired from our previous study using the activated sludge community from the same WWTP under the same incubation conditions.^14^

### 2.2 Cultures and growth conditions

Dechlorinating enrichment culture KB-1 was kindly provided by SiREM lab (https://www.siremlab.com/). *Clostridium homopropionicum* (DSM10017) and *Acetobacterium* species including *A. bakii* (DSM8239), *A. woodii* (DSM1030), *A. fimetarium* (DSM8238), *A. malicum* (DSM4132) *A. wieringae* (DSM1911), *A. paludosum* (DSM8237), and *A. dehalogenans* (DSM11527) were purchased from Deutsche Sammlung von Mikroorganismen und Zellkulturen (DSMZ) as freeze-dried powder. *Acetobacterium* species were revived according to DSMZ protocols in anaerobic glove box filled with N_2_ and H_2_. All cultures were maintained in 160 mL sealed serum bottle containing 95 mL of basal medium prepared according to the recipe on DSMZ website.

### 2.3 Biodefluorination experiment setup

Pure isolates were inoculated into 95 mL sterile basal medium. Fructose and syringic acid were amended according to the recipe on DSMZ website for each strain. 50 µM Cl-PFCA was added as the sole electron acceptor. Aqueous sample was taken periodically during the incubation.

Culture sample was centrifuged at 16,000g for 20 minutes. Supernatant was taken for fluoride measurement and LCMS analysis. Cell pellets were stored properly for DNA (−20 °C) and RNA (−80 °C) extraction, respectively. Abiotic controls were set by adding same amount of Cl-PFCA to sterile medium without inoculation of cultures.

### 2.4 Differential Genes expression evidenced by RNA-sequencing

RNA was extracted using acid-phenol: chloroform: isoamyl alcohol (25: 24: 1) and precipitated in ethanol at – 20 °C as previously described.218 RNA was cleaned up using the RNeasy PowerClean Pro CleanUp Kit (QIAGEN) according to the manufacturer’s instructions. Contaminating DNA in the RNA samples was removed by Turbo DNase Kit (Thermo Fisher Scientific) following the manufacturer’s instructions. The quality of RNA was examined by agarose gel electrophoresis. RNA samples were sent to Microbial Genome Sequencing Center (MiGS) for sequencing analysis. Briefly, RNA samples were treated with RiboZero Plus for rRNA depletion flowed by an Illumina Stranded RNA library preparation procedure and sequenced on a NextSeq 2000 using 2 x 51bp reads.

### 2.5 Crude protein extraction

Cells were collected from 30 mL pre-grown culture by centrifuge culture at 9000 g for 30 min in 2 mL O-ring centrifuge tube with ∼0.1-gram silica beads. For each tube of cells, adding 200 uL 25 mM Tris-HCl buffer (pH 7.5) to re-suspend the cells. Using the cell disruptor to destroy the cell and get the crude enzyme extract with a cycle of 45 seconds on disrupter and cool 45 seconds cooled on ice. The cycle was repeated for 4 times. The lysate was then centrifuged at 10000 g for 20 min. Supernatant was transferred into 5 mL DSMZ *Acetobacterium* medium, CTFE4 was added for degradation examination.

### 2.6 Fluoride measurement

Fluoride ion was measured using an ion-selective electrode (ISE) (HACH). The quantification limit was 0.1 mg/L (c.a. 5.3 µM). The defluorination degree for the removed portion of Cl-PFCA was determined using the formula below:

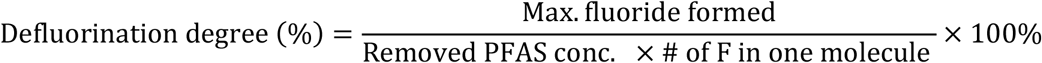

### 2.7 Analytical methods

The parent Cl-PFCA compounds and transformation product suspects were analyzed by an Ultra-high performance liquid chromatography coupled to a high-resolution quadrupole orbitrap mass spectrometer (UHPLC-HRMS/MS, Q Exactive, Thermo Fisher Scientific, Waltham, MA). Two µL sample was injected into a Hypersil Gold column (particle size 1.9 µm, 2.1×100 mm, Thermo Fisher Scientific) and eluted at 0.30 mL/min with water (A) and methanol (B), each containing 10 mM ammonium acetate. The linear gradient was: 95% A for 0 – 1 min, 5% A for 6 – 8 min, and 95% A for 8 – 10 min. Samples were analyzed by a full scan (m/z 50 – 750) at a resolution of 140,000 (m/z 200) under the negative electrospray ionization mode.

### 2.8 Suspect and non-target screening of transformation products

Both suspect screening and nontarget screening were conducted to identify TPs. Briefly, the suspect screening was conducted with Compound Discoverer (Thermo Fisher Scientific).

Plausible TPs were selected based on the following criteria: (i) mass accuracy tolerance < 5 ppm; (ii) isotopic pattern score > 90%; (iii) peak area > 10^5^; (iv)increasing trend along time or first increase and then decrease; (v) no formation in heat-inactivated controls and absent in biological samples without any organofluorine addition; and (vi) not detected as an in-source fragment of the parent compound or any other identified TPs (in-source fragments are peaks identified by an MS full scan, which are also MS^2^ fragments and have the same retention time and formation pattern as the parent compound or the other TPs).

## 3. Results and Discussion

Our previous study discovered that the dechlorinating KB-1 enrichment culture could biodefluorinate PFMeUPA as the solo electron acceptor, and responsible microorganisms belong to minor phylogenetic groups instead of the major dechlorinating species *Dehalococcoides*.^19, 20^ And our recent study revealed that anaerobic activated sludge was capable of defluorinating Cl-PFCAs, which was triggered by dechlorination.^18^ Inspired by these findings, we first used KB-1 enrichment culture to defluorinate CTFE4, which had the highest defluorination degree in activated sludge biotransformation. A near complete defluorination was observed, indicating the Cl-PFCA defluorinating species exist in the KB-1 community (Fig.S1). KB-1 enrichment culture was known to contain acetogen species, however, most of components in the community were yet uncultured without identification of closest relatives at species level. Another recent study from our group reported reductive defluorination of α, β-unsaturated per- and polyfluorocarboxylic acids by *Acetobacterium* species, which are commonly occurring acetogens, revealing the involvement of enzymes of the flavin-based electron-bifurcating caffeate reduction pathway in biodefluorination.^20^ Therefore, we hypothesize that *Acetobacterium* species are capable of biodefluorination for Cl-PFCAs under anaerobic condition. With the observation of defluorination for six Cl-PFCAs, we further investigated the defluorination of CTFE4 with other three *Acetobacterium* species. Fermentation carried out by fermenters was also a key metabolic process occurred in the enrichment culture. *Clostridium homopropionicum* was found to be one of most abundant species in KB-1 culture enriched with PFMeUPA, and it can utilize multiple types of organic substrates. This strain was also examined for CTFE4 defluorination under optimal condition.

### 3.1 Defluorination of and transformation of different Cl-PFCAs by *Acetobacterium bakii*

We investigated ten Cl-PFCAs with different chain length as well as different chlorine substitution degree and position, six of which showed significant fluoride formation (>5 µM) during the 70-day anaerobic incubation compared to the abiotic control (Fig. 1). ∼50 µM PFCAs were fed to *Acetobacterium bakii* to test their biodefluorination under anaerobic condition. No significant parent compound or fluoride formation was observed in abiotic control group (Fig.2 blue symbols). Less transformation products were identified when compared to defluorination carried over by anaerobic activated sludge.^14^ The complicated community structures and resulting versatile metabolic pathways make activated sludge to possess the potential for deeper Cl-PFCA transformation and defluorination.^21^ Interestingly, some Cl-PFCAs could have higher defluorination degree when degraded by pure culture. Interspecies competition may limit the activated sludge to achieve higher defluorination than pure culture for some structures. Another plausible explanation is that the defluorinating species in sludge communities have lower defluorination capacity than the *A. bakii*.

**Figure 2.**
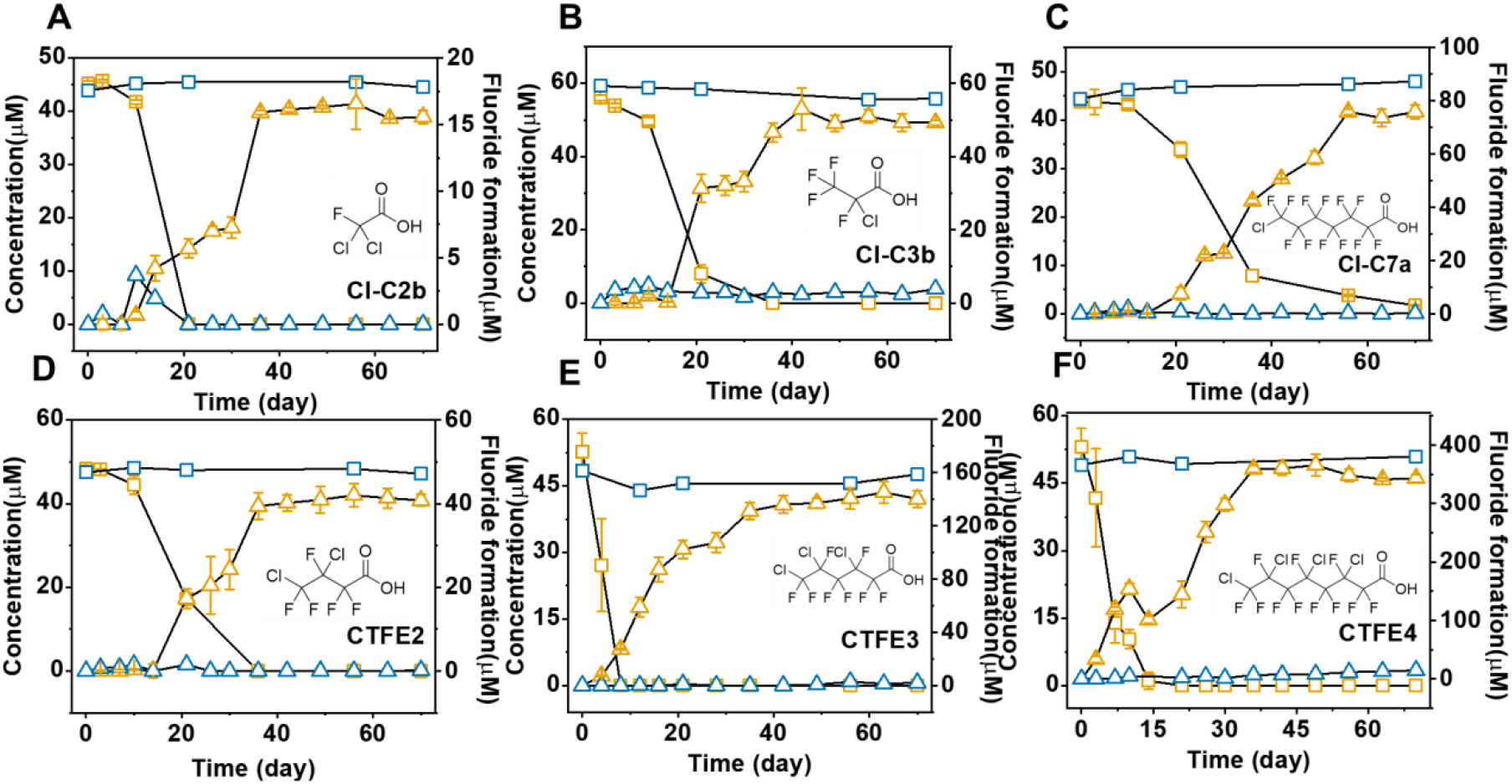
Fluoride formation and parent compound removal of Cl-PFCAs by *A. bakii* in 70-day anaerobic incubation. (A) Cl-C2b, (B) Cl-C3b, (C) Cl-C7a, (D) CTFE2, (E) CTFE3, (F) CTFE4.

Cl-C2b was completely removed within 21 days, however, only 34.5 ± 1.5% fluorine was released as fluoride in biotransformation, indicating existence of non-defluorination transformation products from two third of parent compound (Fig.2A). In comparison, activated sludge can achieve near complete defluorination.^14^ Only one dechlorination transformation product TP110 (─Cl, +H) was identified (Fig.S2A), and its formation matched with the decay of parent compound. The fluoride formation lasted for around 40 days, indicating that at least part of defluorination occurred on intermediates instead of parent compound. Monofluoacetate (MFA) was produced in biodefluorination of Cl-C2b by activated sludge under anaerobic condition,^14^ but not detected in this study. Cl-C2a was not defluorinated in both studies, indicating the one additional chlorine instead of fluorine on alpha carbon enabled the microbial defluorination.

Transformation of Cl-C3b finished within 35 days, and most of fluoride formation came along with the decrease of parent compound (Fig.2B). 22.0 ± 0.6% of fluorine was cut off from Cl-C3b, while only ∼2.5% defluorination was achieved with activated sludge.^14^ TP144 (─Cl, +H) and TP127 (─Cl, ─F, +2H) were identified during the defluorination process both with *A. bakii* and activated sludge. The relative larger peak area of TP144 (∼1×10^8^) over TP127 (∼2×10^6^) indicated major transformation process may stop at the initial dechlorination step. Cl-C3a, which has the chlorine substitution on beta-carbon cannot be defluorinated by pure culture as well as activated sludge.^14^ The result of defluorination of C_2_ and C_3_ Cl-PFCA agreed with the hypothesis that the chlorine substitution at alpha carbon enhanced biodefluorination activity. However, no fluoride formation was observed when Cl-C3c was fed to *A. bakii*.

Cl-C7a had a defluorination degree of 15.4 ± 0.5%, while parent compound was completely transformed in 28 days (Fig.2C). Half of fluoride was released after the consumption of parent compound, indicating that this defluorination may come from further transformation of intermediates generated from parent compound. Major TP was a dechlorination product TP344 (─Cl, +H) (peak area∼2×10^8^), while a dicarboxylic acid TP338 (─Cl, ─F, +H, +2O) was detected with peak areas around 2×10^6^ (Fig.S2F). Interestingly, removal and defluorination of Cl-C7a were both much higher in biotransformation by *A. bakii* than activated sludge, although same TPs were detected in both studies. Cl-5a did not show significant defluorination during the 70-day incubation.

Three CTFE oligomers were tested with *A. bakii* for biodefluorination. Pure culture achieved 16.9 ± 0.3% defluorination for CTFE2, which was higher than the 7% defluorination by activated sludge.^14^ Removal of parent compound and defluorination were both finished in 35 days (Fig.2D). The fluoride was likely to be released via spontaneous defluorination of dechlorination products. Dechlorination product TP210 (─Cl, +H) had largest peak area around 1.6×10^8^. Sequential defluorination product TP174 (─Cl, ─F, +2H) was detected since day 20 and increased to the end of incubation with a peak area of 7×10^6^. TP174 with carbon-carbon double bond was formed via removing one fluorine and one chlorine from parent compound. In addition, a dicarboxylic acid TP170 (─2Cl, ─2F, +2H, +2O) was detected to have a peak area of 6×10^6^ at day 70. TP176 (─Cl, ─F) was only generated by activated sludge and not detected with the pure culture.

Transformation of CTFE3 only took 8 days to complete, while the fluoride was continuously released till day 56, resulting in a defluorination degree of 33.4 ± 4.8% (Fig.2E). In comparison, 20% of organic fluorine was released by activated sludge. Six of seven detected TPs were dechlorination products, while only TP202 (─C, ─3Cl, ─4F, +3H) showed removal of fluorine with a peak area of 1.2×10^5^. TP326 (─Cl, +H), TP292 (─2Cl, +2H), TP259 (─3Cl, +3H) were formed via sequential reductive dechlorination. Carbon-carbon double bond was formed in TP290 (─2Cl) and TP257 (─3Cl, +H). Six additional TPs were discovered in biodefluorination by activated sludge. The biodefluorination by pure culture mainly initiated from reductive (─Cl, +H) and eliminative defluorination (─2Cl), while additional pathways starting from hydrolytic dechlorination (─Cl, +OH) and eliminative defluorination (─HF/─ClF) were proposed for activated sludge.

CTFE4 was removed within 14 days, while fluoride was continuously released till day 49. Biotransformation of CTFE4 resulted in 59.1 ± 5.9% defluorination of total organic fluorine in 70-day incubation (Fig.2F), which was lower than the 77% defluorination achieved with activated sludge. Fourteen of thirty-six TPs detected in biodefluorination with activated sludge were identified in this study, and transformation pathway was proposed based on TP analysis (Fig.3&Fig.S2D). The major CTFE4 transformation pathways were reductive dechlorination and eliminative defluorination (─Cl, ─F), resulting in a series of dechlorination products (i.e., TP442, 408, 374, and 341) and TP422, respectively. In addition, CTFE4 was also transformed to TP406 via eliminative dechlorination (─Cl, ─Cl), which was further dechlorinated and formed TP372.

**Figure 3.**
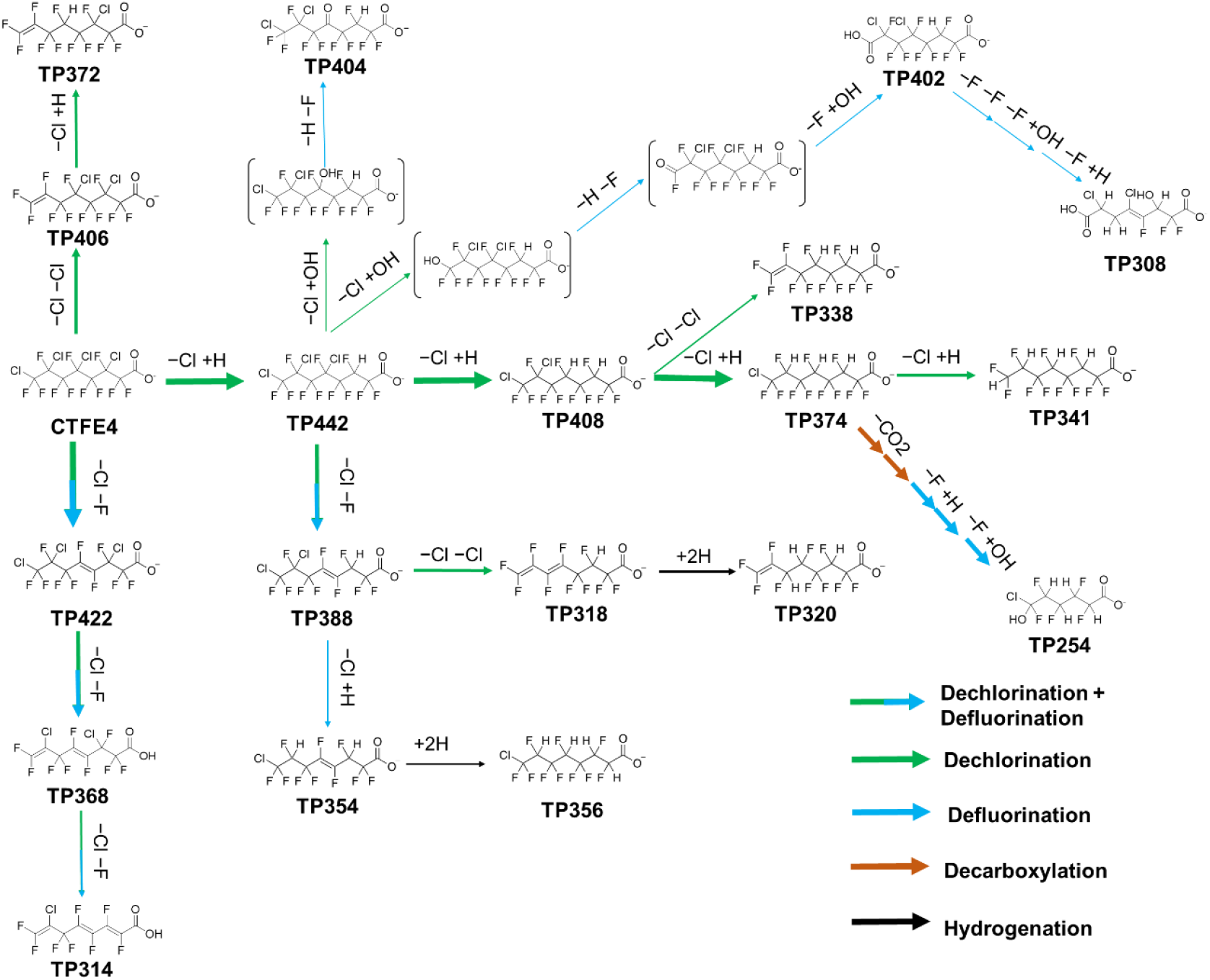
Proposed transformation pathway of CTFE4 biodefluorination by *A. bakii* based on fifteen detected TPs. Green arrow: dechlorination reaction; blue arrow: defluorination reaction, black arrow: hydrogenation reaction; bracket: unstable transient intermediate.

TP388, another major TP, was generated via eliminative defluorination of TP442. Hydrolytic dechlorination and following eliminative defluorination could further transform TP442 to TP404 and TP402, while the position of reaction seemed to be different when generated these two TPs. Chlorines on TP388 and TP408 were further removed via reductive or eliminative dechlorination. TP320 and TP356 had minor formation and were both generated via hydrogenation of carbon-carbon double bond. TPs with m/z lower than 300 were only observed in sludge defluorination experiment, and chain shortening reaction was not observed in biodefluorination by *A. bakii*. Defluorination occurred during chain shortening may lead to the higher defluorination degree in activated sludge.

In summary, the versatile metabolic functions enable activated sludge to transform Cl-PFCAs to additional TPs via different pathway as well as deeper dechlorination and defluorination products than pure culture. However, the defluorination degree could be higher, similar, or lower when Cl-PFCAs were fed to *A. bakii*. The higher defluorination may result from interspecies competition or different responsible enzymes in activated sludge.

### 3.2 Defluorination of CTFE4 by different *Acetobacterium* species

CTFE4 could be biodefluorinated by *A. bakii* and activated sludge with high defluorination degree (>50%). We further investigated its biodefluorination by other three *Acetobacterium* species, as well as *C. homopropionicum* (Fig.4). The defluorination reached the peak between 21-36 days for all *Acetobacterium* species. *A. malicum* achieved 81% defluorination, which was the highest among teste species, and its defluorination was faster than other species. Around 60% defluorination of CTFE4 was observed for the other three species. *A. woodii* and *A. firmetarium* displayed similar defluorination curves, while *A. bakii* took longer time to reach the plateau. This phenomenon was likely to be caused by the reaction kinetics under different growth conditions. All species were maintained under their optimal conditions instructed by DSMZ. Only *A. bakii* was maintained at room temperature, while other species were growing in a 34°C incubator. *C. homopropionicum* was discovered to be one of the most abundant species in KB-1 enrichment culture growing with PFMeUPA as the electron acceptor. However, it did not show any defluorination capacity for CTFE4.

**Figure 4.**
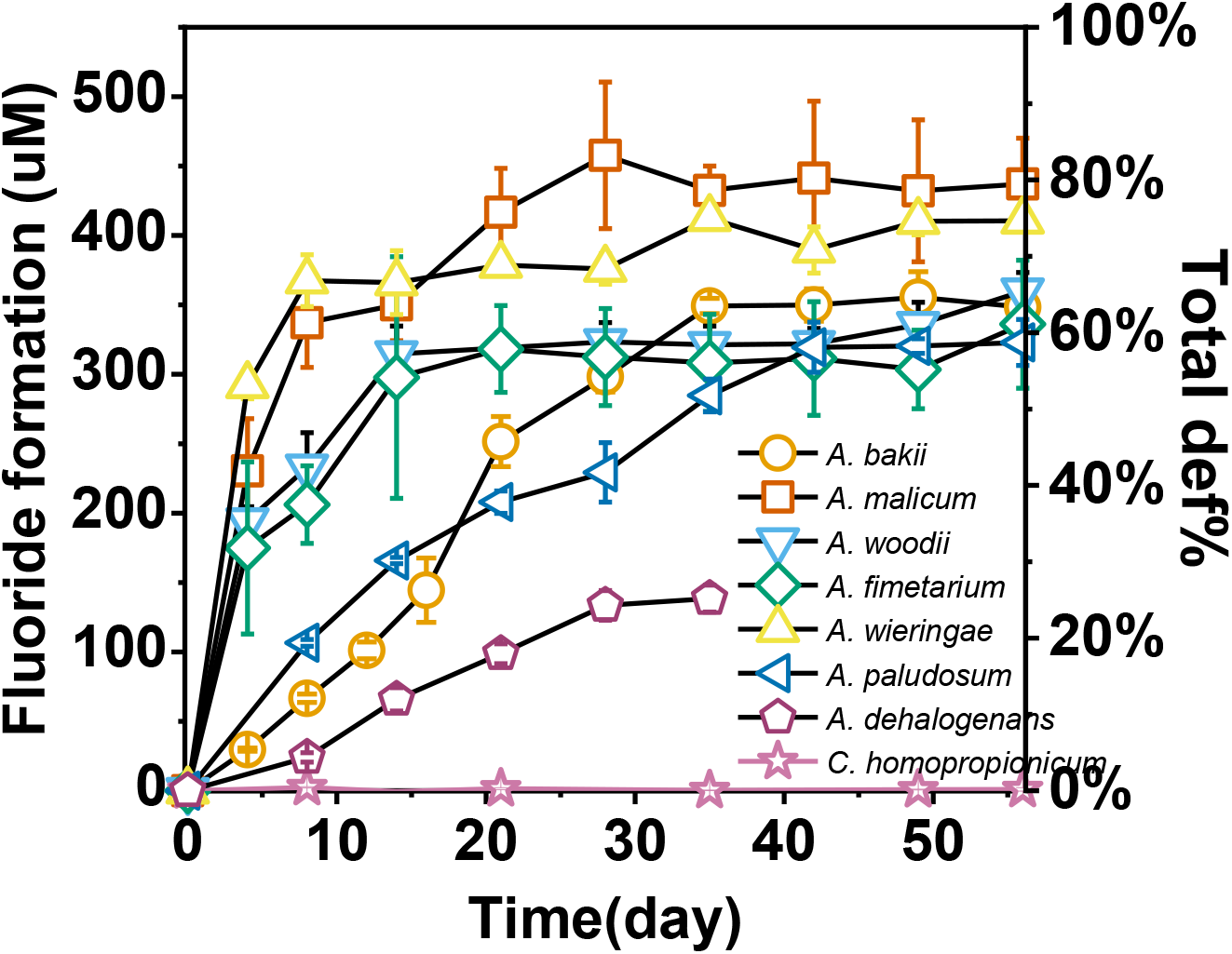
Biodefluorination of 50 μM CTFE4 by *Acetobacterium* species and *C. homopropionicum*.

**Figure 5.**
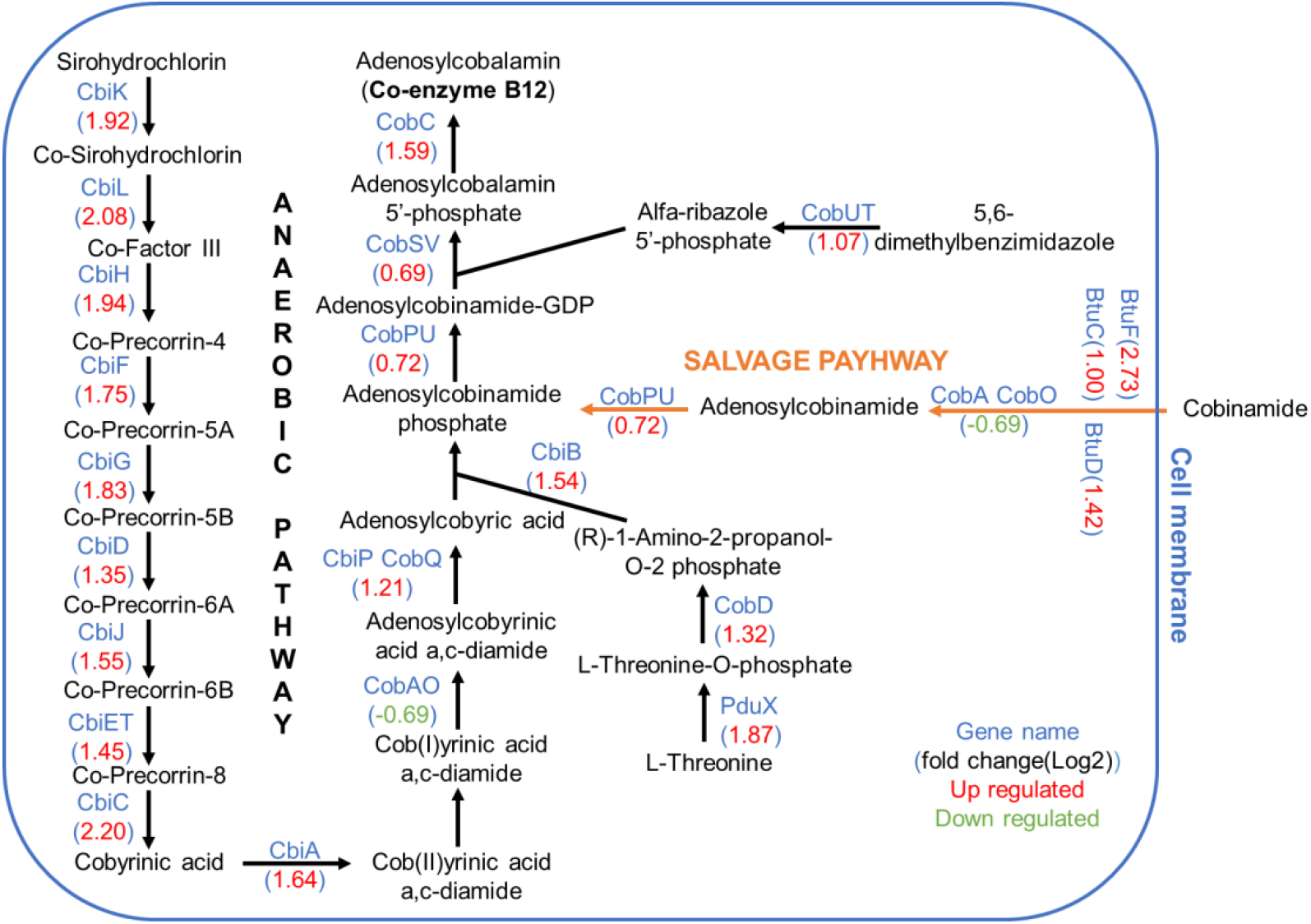
Up regulation of vitamin B12 synthesis related genes with addition of CTFE4 by *A. bakii*. (Note: genes with *p*-value < 0.05, fold-change > 1 were selected: blue dots represent genes with down regulation; red dots represent genes with up-regulation) (n=3).

Crude protein extracted from *A. bakii* under both aerobic and anaerobic conditions were tested for CTFE4 defluorination (Fig.S4). Interestingly, only anaerobically extracted protein could carry out the defluorination, indicating that an oxygen-sensitive enzyme was required to initiate the CTFE4 defluorination. Considering that the protein was extracted from *A. bakii* growing on fructose only, the anaerobically functioning enzymes were possible to be non-specific for dechlorination and defluorination. RNA sequencing was conducted for *A. bakii* growing on fructose only and fructose together with CTFE4.

### 3.3 RNA expression analysis for CTFE4 biodefluorination by *A. bakii*

To further elucidate the reductive defluorination responsible genes and enzymes, we did RNA sequencing for CTFE4 biodefluorination by *A. bakii*. RNA samples were taken in the middle of reductive defluorination processes compared to controls. Genes with significant up/down regulation (i.e., > 1-fold change, *p-value* < 0.05) were selected as suspected genes which regulations were likely related to the CTFE4 biotransformation. The detailed up/down gene regulation are included in Table S2 and Fig. S11. 794 genes were up regulated during CTFE4 biodefluorination, indicating that addition of CTFE4 widely affected the metabolic activities of *A. bakii*. Cobalamins, including vitamin B_12_, are corrinoid-based essential co-factors of reductive dehalogenases. Vitamin B_12_ (100 μg/L) was added to the culture to enhance the anaerobic reductive dehalogenases.^22^ Seventeen cobalamin-related genes were up regulated, including cobalamin synthase and reductase. Sixty-five vitamin B_12_ ATP-binding cassette (ABC) transporters were also found to be up regulated. This result indicated vitamin B12 was likely involved in the CTFE4 dechlorination and defluorination, agreeing with the finding that cobalt-enzyme-dependent anaerobes played a crucial role in the transformation of chlorinated ether PFAS.^15^ The enhanced expression of a chloride channel and a fluoride transport protein were likely to carry out the detoxification by reducing concentrations of the chloride and fluoride.^20, 23^

## 4. Conclusion

Collectively, our results reveal Cl-PFCA biodefluorination by *Acetobacterium* spp. Six of ten selected Cl-PFCAs displayed significant defluorination by *A. bakii* in 70-day anaerobic incubation. Defluorination degrees of Cl-PFCAs by *A. bakii* could be higher or lower than sludge communities, which may attribute to the interspecies competition or diverse metabolic functions in the microbial communities, respectively. Defluorination pathways was proposed and compared with the biodefluorination in sludge communities. Different Acetobacterium species were examined for their capacity on CTFE4 defluorination. Vitamin B12 is likely involved in biodechlorination and biodefluorination of CTFE4, as vitamin B12 synthesis related genes was up-regulated during CTFE4 defluorination. Together, we provide valuable insights on Cl-PFCAs biodefluorination mechanisms.

## Acknowledgment

We would like to give our acknowledgments to SiREM Lab for kindly providing the KB-1 culture. This study is supported by the Strategic Environmental Research and Development Program (ER20-1541 for Y.Y., B.J., C.R., J.L. and Y.M.), the National Science Foundation (Award No. 1931941 for S.C.), National Institute of Environmental Health Sciences (Award No. 1R01ES032668-01 for S.C.).

## Notes

### Competing Interest Statement

The authors have declared no competing interest.

