## Supplementary material for "Biotransformation and biodefluorination of chlorinated polyfluorocarboxylic acids by *Acetobacterium* species": SI

[2] State Key Laboratory of Chemical Safety, SINOPEC Research Institute of Safety Engineering Co., Ltd., Qingdao 266071, China (current address)

[3] Department of Civil and Environmental Engineering, Stanford University, 473 Via Ortega, Stanford, CA 94305, United States (current address)

**Corresponding Author**

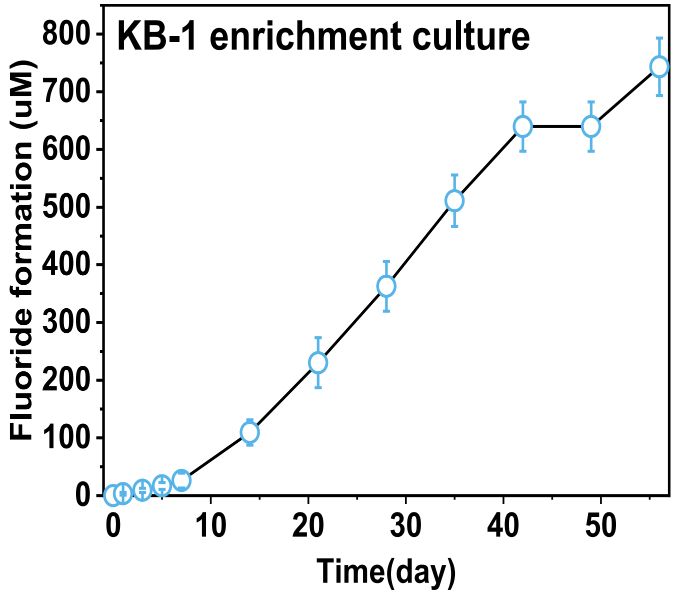


Figure S1. Fluoride formation of CTFE4 biodefluorination by KB-1 enrichment culture. ~80 μM CTFE4 was almost completely defluorinated with defluorination degree over 80%.

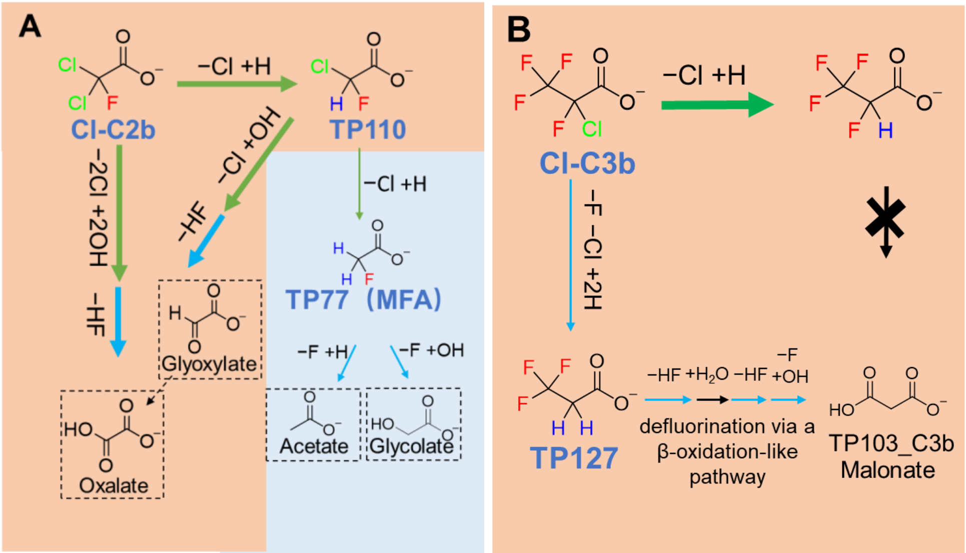

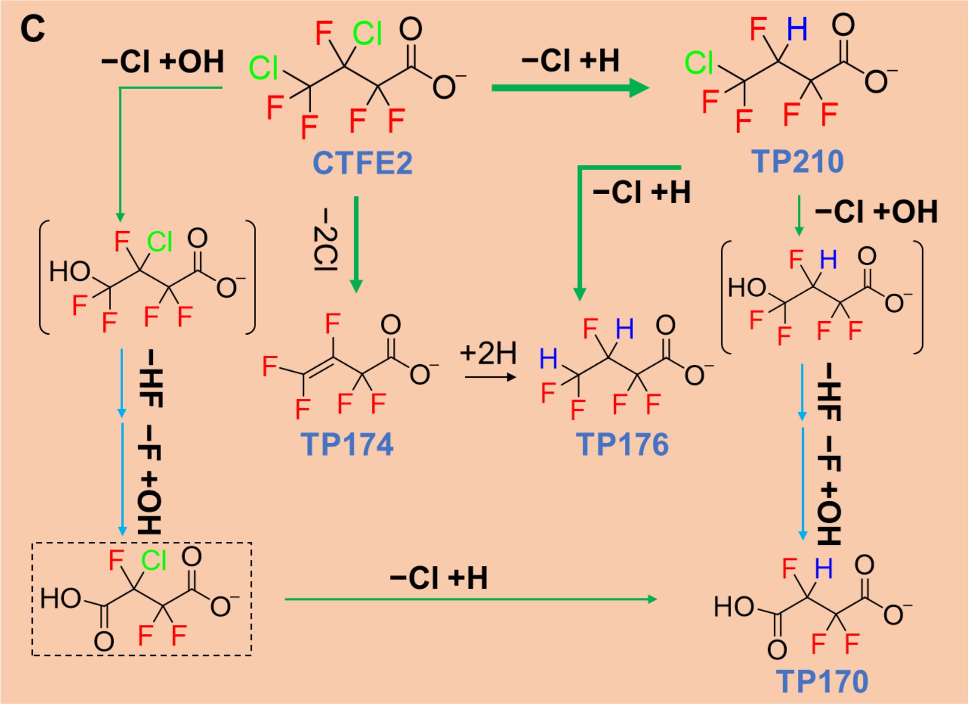
C
−Cl +OH
CTFE2
−Cl +H
TP210
−2Cl
−Cl +H
−Cl +OH
+2H
TP174
TP176
−HF
−F +OH
−HF
−F +OH
−Cl +H
TP170

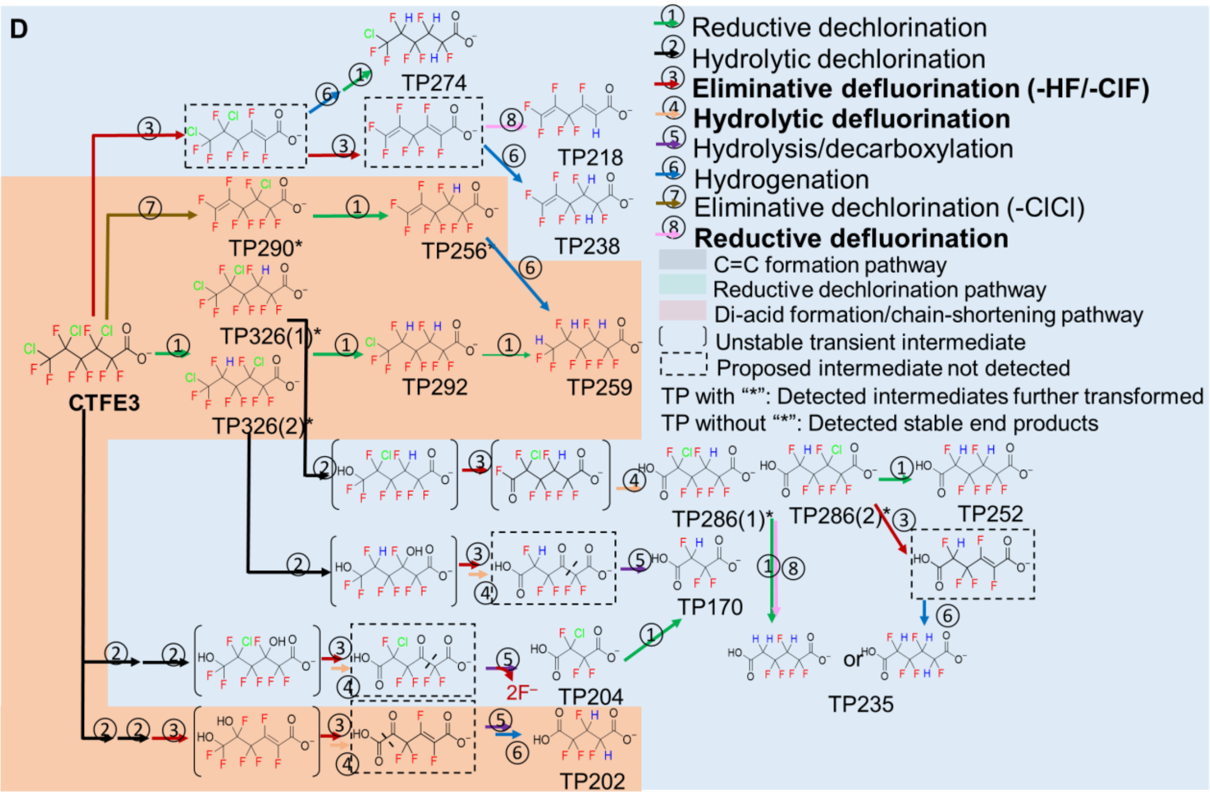
D
TP274
TP218
TP290*
TP256*
TP238
TP326(1)*
TP326(2)*
TP292
TP259
CTFE3
TP286(1)*
TP286(2)*
TP252
TP170
TP235
TP204
TP202
2F⁻
or
① Reductive dechlorination
② Hydrolytic dechlorination
③ Eliminative defluorination (-HF/-ClF)
④ Hydrolytic defluorination
⑤ Hydrolysis/decarboxylation
⑥ Hydrogenation
⑦ Eliminative dechlorination (-ClCl)
⑧ Reductive defluorination
C=C formation pathway
Reductive dechlorination pathway
Di-acid formation/chain-shortening pathway
Unstable transient intermediate
Proposed intermediate not detected
TP with "*": Detected intermediates further transformed
TP without "*": Detected stable end products

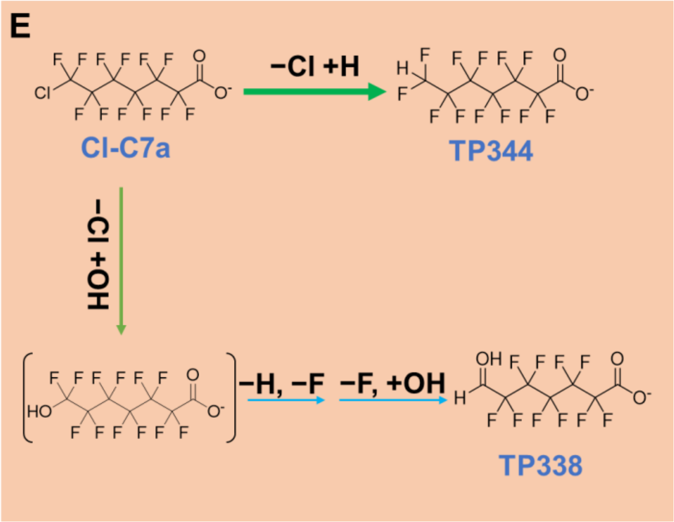


Figure S2. Comparison of Cl-PFCAs transformation pathways between *A.bakii* and activated sludge. (A) Cl-C2b; (B) Cl-C3b; (C) CTFE2; (D) CTFE3; (E) Cl-C7a. Orange color represents the pathways observed in *A. bakii* defluorination experiment.

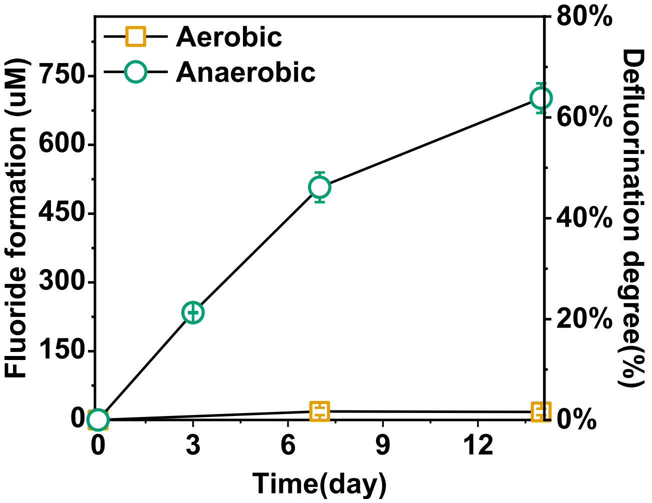


Figure S3. Defluorination of CTFE4 by crude protein extracted from *A. bakii* under aerobic and anaerobic condition.

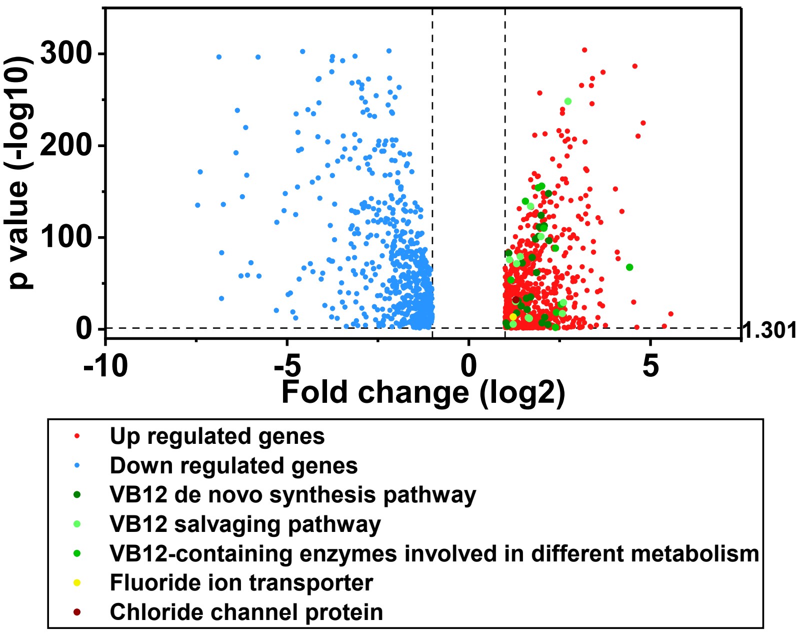


Figure S4. Differential gene expression in *A. bakii* growing with CTFE4.

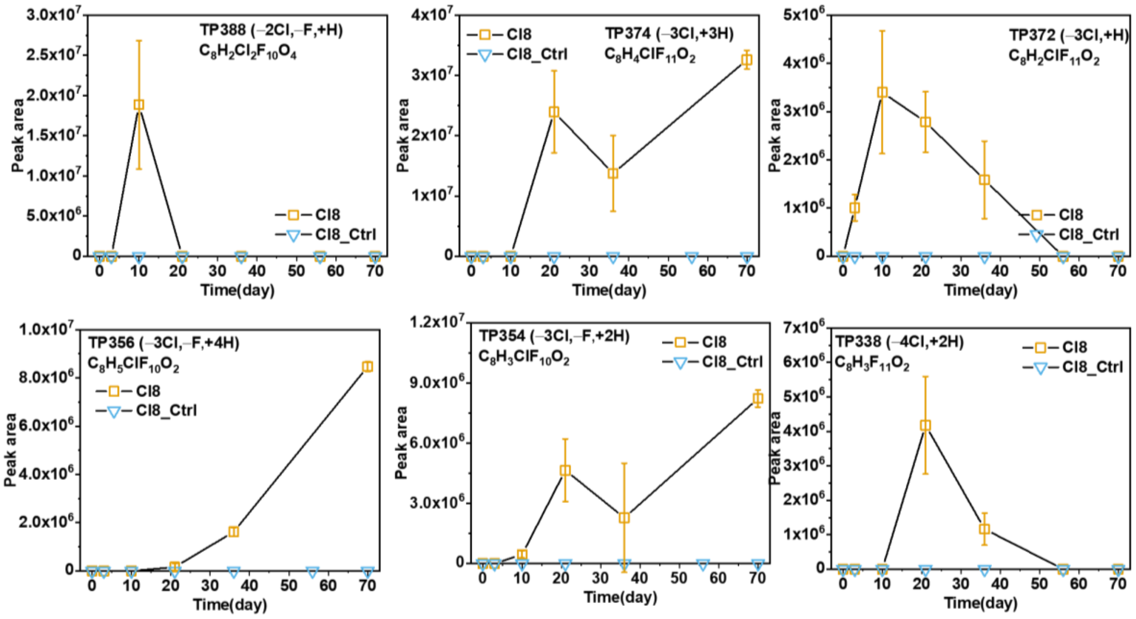

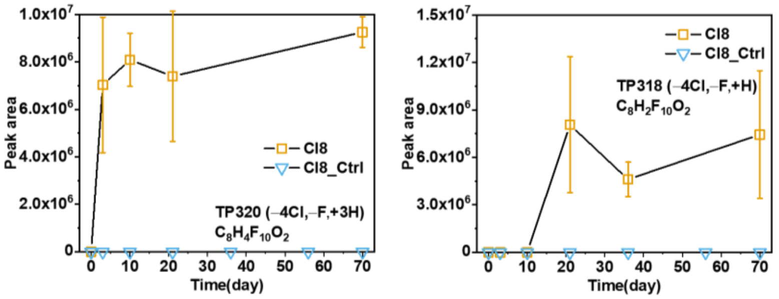


Figure S5. TPs of CTFE4 biodefluorination by *A. bakii*.

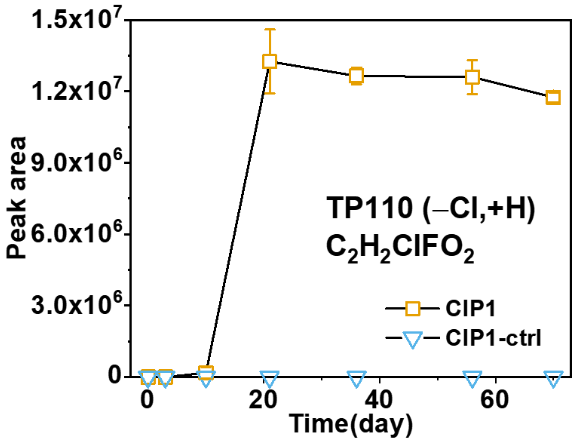


Figure S6. TP of Cl-C2b biodefluorination by *A. bakii*.

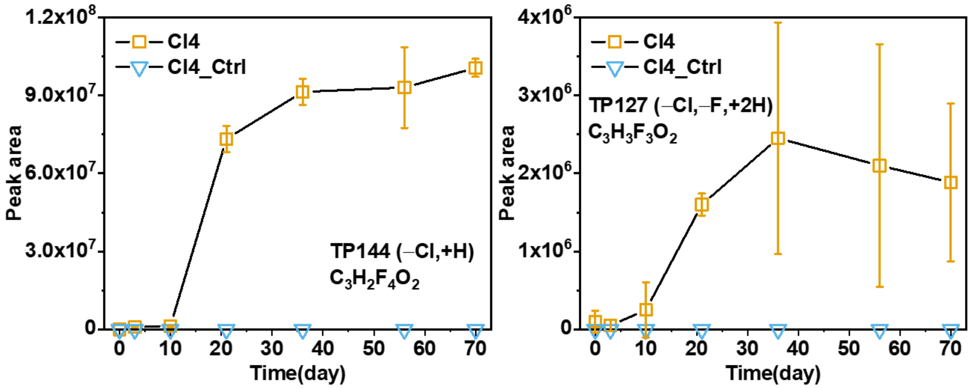


Figure S7. TPs of Cl-C3b biodefluorination by *A. bakii*.

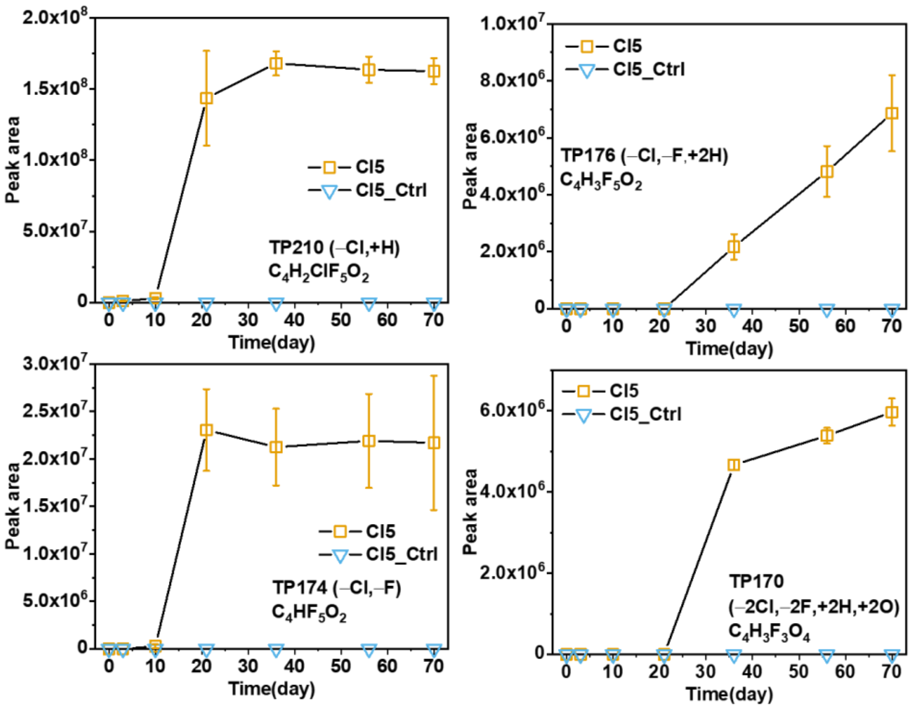


Figure S8. TPs of CTFE2 biodefluorination by *A. bakii*.

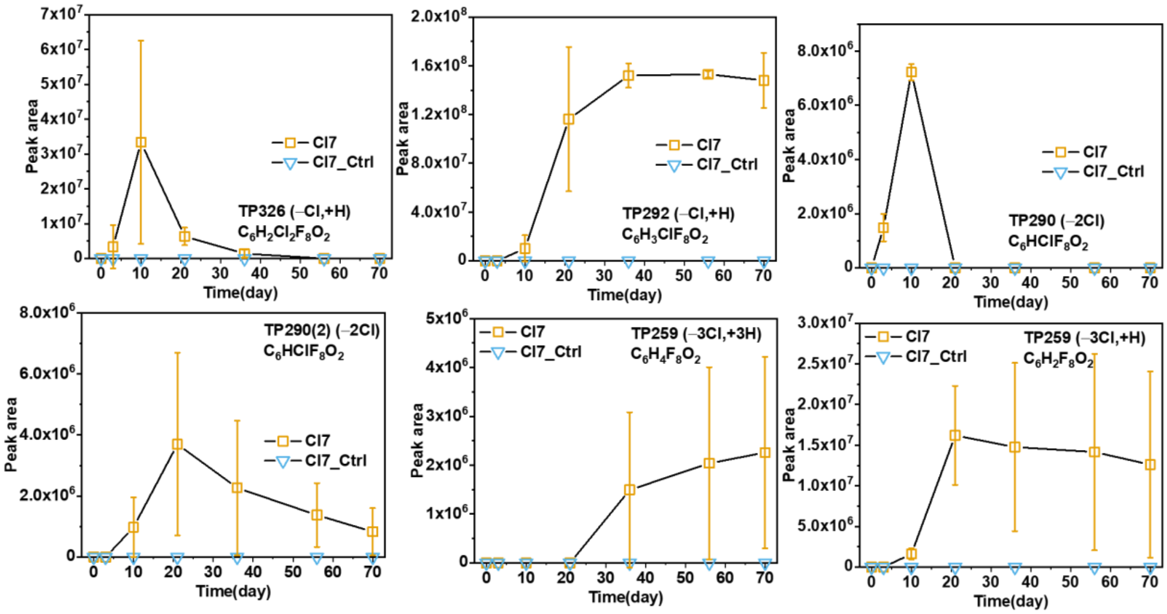


Figure S9. TPs of CTFE3 biodefluorination by *A. bakii*.

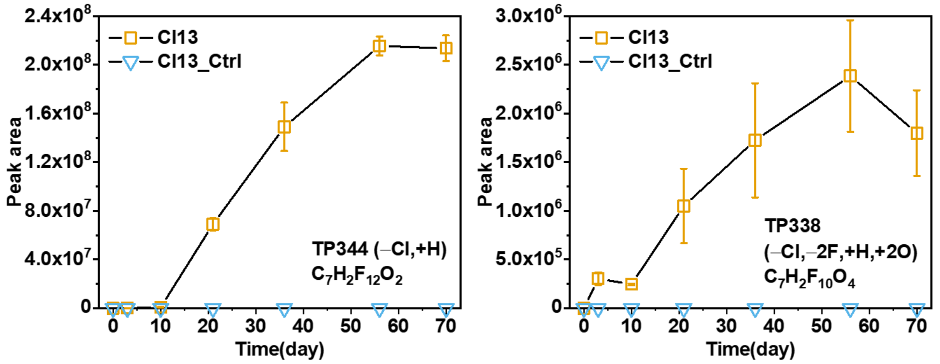


Figure S10. TPs of Cl-C7a biodefluorination by *A. bakii*.

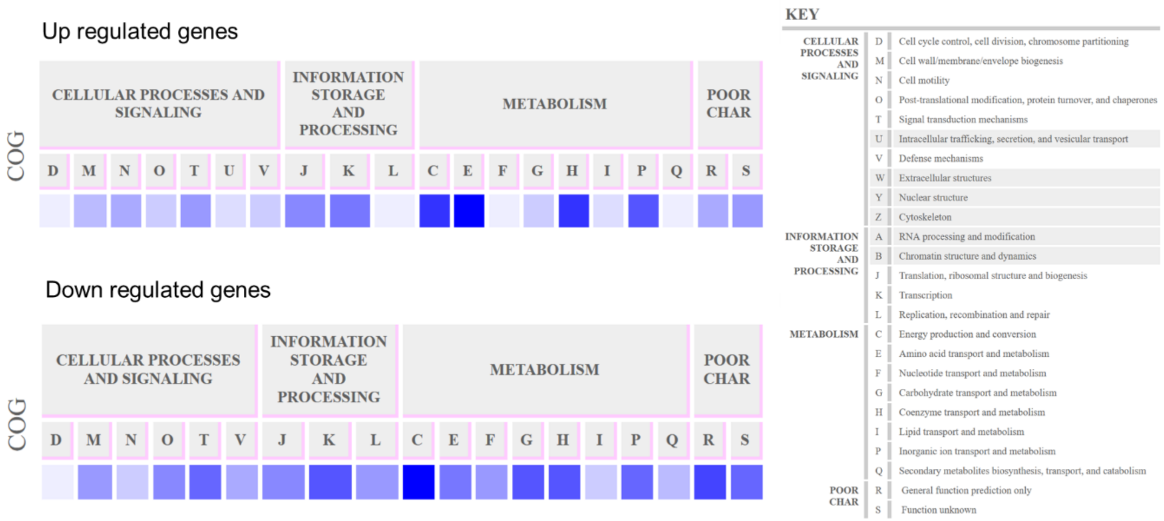


Figure S11. Differential gene expression during biotransformation of CTFE4.
